# Altered cellular VEGF dynamics in Chronic Obstructive Pulmonary Disease

**DOI:** 10.64898/2026.05.04.722512

**Authors:** Maria del Pilar Romano, Pia Ecke, Ellen Tufvesson, Sukhwinder Singh Sohal, Leif Bjermer, Martina Schmidt, Gunilla Westergren-Thorsson, Anna-Karin Larsson-Callerfelt

## Abstract

Pulmonary vascular remodelling is common in patients with chronic obstructive pulmonary disease (COPD). Vascular endothelial growth factors (VEGFs) are key mediators in angiogenesis and vascular remodelling and exist in different isoforms. VEGF-A is the most potent angiogenic member binding to VEGF receptor 2 (VEGFR2). There are, however, few studies on other isoforms, as VEGF-C, and its receptor VEGFR3 in COPD and subsequent impact of cAMP therapies on VEGF isoforms. Our aim was to evaluate the VEGF isoform synthesis in primary distal lung fibroblasts from control subjects (non-smokers (n=6) and ex-smokers (n=4), and COPD subjects with GOLD stage II (n=4) or GOLD stage IV (n=6), and the expression of VEGFR2 and VEGFR3 in human lung tissue. Primary lung fibroblasts were exposed to the cAMP generating therapies formoterol, iloprost, or roflumilast, the adenylyl cyclase activator forskolin or to transforming growth factor (TGF)-b1. VEGF isoforms were evaluated with ELISA. VEGF-C release was not significantly altered by TGF-β1, in contrast to the increased levels of VEGF-A, in all fibroblasts. VEGF-C was significantly decreased by iloprost, forskolin and formoterol, whereas VEGF-A was significantly increased by iloprost and forskolin, with differences in release pattern between and within fibroblasts from control and COPD subjects. Exposure to VEGF-C specifically towards VEGFR3 decreased proliferative rate in human lung fibroblasts and bronchial epithelial cells. VEGFR2 and VEGFR3 were both present in parenchymal lung tissue and VEGFR2 in pulmonary blood vessels. in both healthy and COPD, whereas there was elevated expression of VEGFR3 in bronchial epithelium. In conclusion, TGF-β1 and cAMP generating compounds have significant effects on VEGF-C and VEGF-A synthesis, which appear dysregulated in lung fibroblasts from ex-smokers and patients with COPD. Increased VEGFR3 expression in the bronchial epithelium in lung tissue, and studies into their functional impact, warrants further investigations.

## Introduction

Chronic Obstructive Pulmonary Disease (COPD) is a severe heterogeneous lung disease with no cures or effective treatments available. The disease is characterised by airflow limitations linked to chronic inflammation, fibrosis in small airways, and emphysema due to destruction of alveolar epithelial cells and capillaries (1). Lack of sufficient oxygen supply in the lung results in hypoxic milieus which further trigger cellular activity and pathological remodelling processes in airway (2). These events contribute to decline in effective oxygen exchange in the alveoli and to progression of the disease with systemic complications and cardiovascular comorbidities (3). Hypoxic events induce vascular endothelial factor (VEGF) to increase oxygen delivery by promoting angiogenesis but also vascular remodelling (2). In COPD, pulmonary vascular remodelling is common (3) and comorbidities in cardiovascular disease have negative impacts on COPD prognosis (4). Increased VEGF expression has been associated with bronchial angiogenesis that inversely correlate with lung function in COPD patients (5). VEGF exists in different isoforms where VEGF-A isoform 165 is the most potent angiogenic member binding to VEGF receptor 2 (VEGFR2) (6). Another isoform is VEGF-C which is linked to lymphangiogenesis via activation of VEGFR3 (7-9) and promotes angiogenesis via VEGFR2/VEGFR3 complex (10). VEGF-C expression is primarily related to be induced during hypoxia in tumours (11) whereas limited data is available about the role of VEGF-C in the healthy lung and COPD pathology. Fibroblasts are key players in regulating the homeostasis of extracellular matrix (ECM) but also in pathological remodelling processes by constituting a rich source of cytokines and growth factors in cross talks with other structural cells and immune cells (12). We have previously shown that fibroblasts from COPD patients have an altered phenotype (13-15). Our previous data show that both VEGF-A and VEGF-C is synthesised by primary human lung fibroblasts ((13, 14) and that VEGF-A is induced by transforming growth factor (TGF)-b (14). We have previously shown that VEGF-A release is induced by the prostacyclin analogue iloprost (14). Prostacyclin analogues, b_2_ agonists, as formoterol, and phosphodiesterase-4 (PDE_4_) inhibitors, as roflumilast, are cAMP generating drugs (16), and both formoterol and roflumilast are commonly administered to patients with COPD. The anti-fibrotic effect of cAMP generating therapies is partly related to decrease in collagen I and III synthesis (15-19). The effect of cAMP generating therapies on VEGF synthesis is however less known. VEGF may induce different effects depending on which isoforms that are synthesised in lung fibroblasts and which receptor that is expressed. The aim of this study was therefore to further investigate the effects of cAMP generating drug therapies on lung fibroblasts and the expression of VEGF receptor in lung tissue obtained from control subjects and COPD patients. Our obtained data indicate that TGF-β1 and cAMP generating compounds have different effects on VEGF-A and VEGF-C synthesis and that the response is altered between non-smokers, ex-smokers and COPD patients. VEGFR2 and VEGFR3 are expressed at different localizations in the lung. The expression of VEGFR3 in the bronchial epithelium warrants further investigations.

## Methods

### Patient characteristics

Primary lung fibroblasts and lung tissue were obtained from subjects with COPD undergoing lung transplantation or tumour resection at Skåne University hospital or lung transplantation at Sahlgrenska University hospital, Gothenburg, Sweden and from healthy lung donors. All COPD patients were ex-smokers who stopped smoking six months before the transplantation or tumour resection. Lung explants from healthy organ donors (n = 8), with no history of cigarette smoking or lung disease were included in the study (Table 1). The donor lungs were intended for transplantation but approved for research when the organ did not meet clinical grade upon arrival or when matched recipients were lacking. Written consent was obtained from their closest relatives. Tissues from ex-smokers (n = 4) and COPD GOLD stage 2 patients (n = 4) were collected from tumour resections performed at Lund University hospital. Lung tissue that was used was taken as far away from the tumour as possible and was evaluated to be macroscopically healthy by an experienced pathologist. In addition, tissue was histologically assessed by haematoxylin and eosin (H&E) staining to ensure the absence of infiltrating cancer cells. Tissue from GOLD stage IV patients (n = 9) were from lung explants from transplantations performed at Sahlgrenska University hospital in Gothenburg. All COPD donors provided written informed consent, and the study was performed in accordance with the Declaration of Helsinki. COPD patients had a postbronchodilator FEV1/FVC ratio of less than 0.7 and the healthy smokers had normal spirometry with FEV1/FVC ratio > 0.7 (Table 1). Ethical approval for the study was obtained from the Swedish Ethical Review Authority (ethical permit numbers: FEK 91/2006, FEK 413/2008, 675-12-2012, 1026-15, 2015-120), and all experiments complied with the relevant guidelines. All patients were decoded such that they could not be identified in our study.

**Table 1.** with patient characteristics. ^*^Lung tissue obtained after resection.

| Subject | Lung disease | Smoking status | Pack years | Sex | Age/age range | DLCO (%) | FEV1 | FVC | FEV1/FVC (%) |
| --- | --- | --- | --- | --- | --- | --- | --- | --- | --- |
| 1 | Healthy | never | N/A | Male | 48 | N/A | N/A | N/A | N/A |
| 2 | Healthy | never | N/A | Male | 31-45 | N/A | N/A | N/A | N/A |
| 3 | Healthy | never | N/A | Male | 62 | N/A | N/A | N/A | N/A |
| 4 | Healthy | former | N/A | Male | 66 | N/A | N/A | N/A | N/A |
| 5 | Healthy | never | N/A | Female | 41 | N/A | N/A | N/A | N/A |
| 6 | Healthy | never | N/A | Male | 56 | N/A | N/A | N/A | N/A |
| 7 | Healthy | never | N/A | Male | 39 | N/A | N/A | N/A | N/A |
| 8 | Healthy | former | N/A | Male | 62 | N/A | N/A | N/A | N/A |
| 9 | Healthy* | former | 30 | Female | 69 | 50 | 2.0 | N/A | N/A |
| 10 | Healthy* | former | 18 | Female | 36 | 64 | 1.78 | N/A | N/A |
| 11 | Healthy* | former | 20 | Female | 65 | 96 | 2.5 | N/A | N/A |
| 12 | Healthy* | former | 25 | Male | 71 | 62 | 2.69 | N/A | N/A |
| 13 | COPD GOLD 2* | former | 31 | Female | 68 | 91 | 2.27 | N/A | N/A |
| 14 | COPD GOLD 2* | former | 51 | Male | 67 | 56 | 2.18 | N/A | N/A |
| 15 | COPD GOLD 2* | former | N/A | Male | 66 | N/A | N/A | N/A | N/A |
| 16 | COPD GOLD 2* | former | 41 | Female | 73 | 81 | 1.54 | N/A | N/A |
| 17 | COPD GOLD 2* | former | 30 | Female | 66 | 27 | 1.27 | 2.85 | 58 |
| 18 | COPD GOLD 4 | former | 40 | Male | 59 | N/A | 0.5 | 1.3 | N/A |
| 19 | COPD GOLD 4 | former | N/A | Female | 59 | 33 | 0.4 | 1.3 | 40 |
| 20 | COPD GOLD 4 | former | 35 | Female | 63 | 45 | 0.6 | 1.9 | 40 |
| 21 | COPD GOLD 4 | Former | 60 | Male | 59 | 16 | 1 | 2.69 | 50 |
| 22 | COPD GOLD 4 | former | N/A | Female | 69 | 29 | 0.5 | 1.6 | 39 |
| 23 | COPD GOLD 4 | former | 4.6 | Female | 61 | 38 | 0.4 | 1.5 | 22 |

### Culture of primary distal lung fibroblasts

Primary distally-derived lung fibroblasts from the lung tissue sections were cultured in Dulbecco’s Modified Eagle Medium (DMEM, Sigma-Aldrich, St Louis, MO USA) with 10% foetal clone serum (FCIII, Thermo Fisher scientific, Waltham, MA, USA), 1% Gentamicin, 1% Amphotericin and 1% L-glutamine (all from Gibco BRL, Paisley, UK), as previously described (14). Cells were cultured at 37°C with 10% CO_2_. The primary distal lung fibroblasts were used in passage 4-7. Before stimulation the medium was changed to 0.4% FCIII with 1% Gentamicin, 1% Amphotericin and 1% L-glutamine to avoid pre-existing growth factors in the serum. Exposures to either iloprost (1 µM, Cayman Chemical, Ann Arbour, US), forskolin (1 µM, Sigma-Aldrich), roflumilast (1 µM) (Cayman Chemical, formoterol (1 µM, Sigma-Aldrich), or TGF-b1 (10 ng/mL, #240-B, RD Systems) and a unstimulated control, containing only medium with 0.4% serum. The cells were exposed for 24 h at 37°C with 10% CO_2_ before they were harvested. Supernatants from all wells were collected and stored at −80 °C. The cells were harvested using 100 µL RIPA lysate buffer mixed with cOmplete protease inhibitor cocktail (Sigma-Aldrich, St Louis, MO USA) and centrifuged for 10 min at 4°C with 14,000 rpm. The protein supernatant was collected and stored at −80°C.

### Enzyme-linked immunosorbent assay for analysis of VEGF-A, C and D

VEGF-A, VEGF-C and VEGF-D were measured using Human VEGF Quantikine ELISA kit (DVE00, DVEC00, DVED00, R&D Systems, Abingdon, UK, and the manufacturers protocols were followed. Absorbance was measured at 450 nm using a microplate reader (Multiskan GO, Thermo Fisher Scientific). The assay detection limit for VEGF-A was 15.6 pg/mL, 108 pg/mL for VEGF-C and 125 pg/mL for VEGF-D.

### Immunofluorescence staining

Staining of lung tissue from healthy donors, GOLD 2 and GOLD 4 COPD patients was selected based on the presence of one or more airways. The tissue was sectioned at 4 µm thickness using a microtome and stained with hematoxylin and eosin (HE) to identify and validate the presence of airways. Parallel tissue slides were stained for VEGFR3. The slides were deparaffinized and antigen retrieval were performed at low pH 6.1. Slides were then washed 3 x 5 min in tris buffered saline (TBS) and stained overnight with a primary rabbit anti-human VEGFR3 antibody (ab27278, Abcam, Cambridge, United Kingdom) diluted 1:100 in dilution buffer (TBS + 2 % bovine serum albumin (BSA)). Following 2 x 5 min wash in TBS, slides were incubated for 45 min with secondary donkey anti-rabbit IgG Alexa fluor 488 conjugate antibody (21206, Thermo Fisher Scientific) diluted 1:200 in dilution buffer, supplemented with 4′,6-Diamidino-2-phenylindole dihydrochloride (Dapi) for nuclear staining. Following a final 2 x 5 min wash, slides were mounted with fluorescence mounting medium (Dako) and scanned in an Olympus Automatic Virtual Slide Scanner System VS-120, with the exposure times 16 and 200 ms for the DAPI and FITC channel respectively. An image of every airway in every section was captured in the Olympus OlyVia software. VEGFR3 and pan-cytokeratin co-staining were made on parallel tissue sections from nine of the subjects included in the VEGFR3 staining (n=3 per group). The procedure was the same except for the addition of a primary mouse anti-human pan-cytokeratin AE1/AE3 antibody (ab27988, Abcam), diluted 1:100.

### Cell culturing of human lung fibroblasts and bronchial epithelial cells

The cell lines human bronchial epithelial cells BEAS2-2B and human lung fibroblasts (HFL-1) were used to evaluate the effect of VEGF-C on cell proliferation and expression of VEGFR3. BEAS-2B were used in passage 29 to p31 and HFL-1 cells in passage 16 to 19. The cells were seeded (7000 cells/well) in 96-well plates (Sarstedt) in RPMI 1640 and DMEM medium, respectively (61870, 61965-026, Gibco) supplemented with 10% FCII serum and 1% AB/AM for 24 h. The medium was then changed and the cells were either stimulated with 0.1 ng/mL, 1 ng/mL, 10 ng/mL and 100 ng/mL of Recombinant Human VEGF-C (9199-VC, RD Systems) (VEGF-C ligand binding to both VEGFR2 and VEGFR3) or Recombinant Human VEGF-C (Cys156Ser) (752-VC, RD Systems) (specific VEGF-C ligand for VEGFR3) for 24, 48 and 72 h. Proliferation rate was measured as previously described (14). Briefly, cells were fixed in 1% glutaraldehyde in PBS (18912014, Gibco™) for 30 min, washed with PBS and then stained with 0.1% crystal violet (HT90132, Sigma-Aldrich). The plates were then washed with water to remove excessive staining. 200 mL of 1% Triton™ X-100 (T9284, Sigma-Aldrich) was added to each well and incubated overnight. The absorbance was read at 595 nm with a spectrophotometer plate reader. Effect of VEGF-C was expressed as fold change compared to control. To evaluate VEGFR3 expression and release of VEGF-C, BEAS-2B and HFL-1 cells were seeded (2 × 10^5^ cells/well) in 6-well plates (Nunc, Sarstedt) and cultured until 80 % confluency in RPMI 1640 and DMEM medium, respectively), supplemented with 10% FCIII serum and 1% AB/AM. Medium was changed to RPMI or DMEM medium with 1% serum for 2 h before the cells were exposed to 10 ng/mL of Recombinant Human TGF-beta 1 (#240-B, RD Systems) or 1 ng/mL of 10 ng/mL of either Recombinant Human VEGF-C (9199-VC, RD Systems) or Recombinant Human VEGF-C (Cys156Ser) (752-VC, RD Systems) for 48 h. Total protein was collected from BEAS-2B and HFL-1 lysed with RIPA Lysis Extraction Buffer (89900, Thermo Scientific™) supplemented with cOmplete™, EDTA-free Protease Inhibitor Cocktail (COEDTAF-RO, Roche) and PhosSTOP™ (PHOSS-RO, Roche). Cell homogenates were centrifuged at 14000 g for 30 min at 4°C, and the supernatant was collected as total protein extract.

### Total protein concentration in cell lysate

The total amount of proteins in the lysate from the primary lung fibroblasts, HFL-1 and BEAS-2B was analysed using Pierce BCA Protein Assay Kit (23225, Thermo Scientific). A bovine serum albumin (BSA) standard was made ranging from 2000 µg/ml to 62.5 µg/ml and a blank. Absorbance was measured at 562 nm with a microplate reader (Multiskan GO, Thermo Fisher Scientific).

### Western Blot

To verify VEGFR3 expression on the fluorescence staining, we performed protein expression of VEGFR3 on human lung fibroblasts and human bronchial epithelial cells. The samples were heated at 95 °C for 20 min and 15 µg of protein was run on a 4–15% Mini-PROTEAN TGX Stain-Free Protein Gels (#4568086, Bio-Rad) using 4x Laemmli Sample Buffer (#1610747, Bio-Rad). Electrophoresis was performed at 180 V for 1 h, followed by Stain-Free gel activation for 45 sec using a ChemiDoc Imaging System. Following the gel activation, proteins were transferred on the membranes of the Trans-Blot Turbo RTA Mini 0.2 µm Nitrocellulose Transfer Kit (#1704270, Bio-Rad) using Trans-Blot® Turbo™ Transfer System (#1704150, Bio-Rad) 1.3A, 200V for 10 min. The stain-free blot image is used for total protein loading control and normalization. Membranes were blocked with EveryBlot Blocking Buffer (#12010020, Bio-Rad) for 20 min and then probed overnight at 4°C with Rabbit Recombinant Monoclonal VEGF Receptor 3 antibody (ab243232, Abcam) diluted 1:1500 in 5% BSA in TTBS (10x Tris Buffered Saline, #1706435, Biorad, 0,1% TWEEN® 20, P1379, Sigma-Aldrich). Goat Anti-Rabbit IgG H&L (HRP) (ab97051, Abcam) diluted 1:10 000 in 2% BSA in TTBS, was added at room temperature for 1 h. Immune-reactive proteins were detected using the Clarity Max Western ECL Substrate (#1705062, Bio-Rad) according to the manufacturer’s instructions. Densitometric analysis was performed on the ∼75 kDa VEGFR3-immunoreactive band. Quantification was performed only on this band, as other faint bands were inconsistently observed and not reliably measurable. Protein levels were normalized to total protein signal of the stain free blot and expressed as fold change relative to untreated control samples within each independent experiment. All experiments were performed in quadruplicate. The amount of protein per lane for Stain-Free images was quantified using the lane and bands tools in Image Lab v6.1 (Bio-Rad). All calculations were performed automatically by the software.

### Statistical analysis

Data are shown for individual subjects as absolute values and presented as median. The non-parametric Mann–Whitney t-test was used to compare statistical differences between two groups. Two-ways repeated measurement analysis of variance (ANOVA) on ranks followed by the non-parametric Dunn’s post hoc test were used to compare differences in pharmacological treatments.

Data for BEAS-2B and HFL-1 are presented as fold change to control with mean and SEM and statistical analysis were performed with one-sample t-test. Differences were considered to be statistically significant at P < 0.05. All analyses were performed using GraphPad Prism 10.5.0 (GraphPad Software Inc., San Diego, CA, US).

## Results

### Release of VEGF-C compared to VEGF-A in primary lung fibroblasts

The release of VEGF-C was significantly lower in lung fibroblasts from COPD patients (n=10) compared to control subjects (n=10) at baseline (unstimulated conditions) (p=0.0094) (fig 1). When subgrouped, COPD GOLD 2 (n=4) and GOLD 4 (n=6) showed both a significantly lower release of VEGF-C (p<0.05) compared to former smoker (n=4) (fig 1.A-D). There were no differences between ex-smokers and non-smokers (n=6). Stimuli with TGF-b 10 ng/mL did not significantly alter the release of VEGF-C in either lung fibroblasts from control subjects or patients with COPD (Fig 1.A-D). Treatment with different cAMP mediating compounds showed that forskolin and iloprost significantly decreased the release of VEGF-C in fibroblasts from healthy lung donors (fig 1.A) and ex-smokers (fig. 1.B), whereas forskolin significantly reduced VEGF-C release in COPD GOLD 2 (fig 1.C) and GOLD 4 (fig 1.D). There was a trend towards reduced VEGF-C levels to iloprost in COPD GOLD 2 (p=0.059) (Fig 1.C).

**Figure 1.**
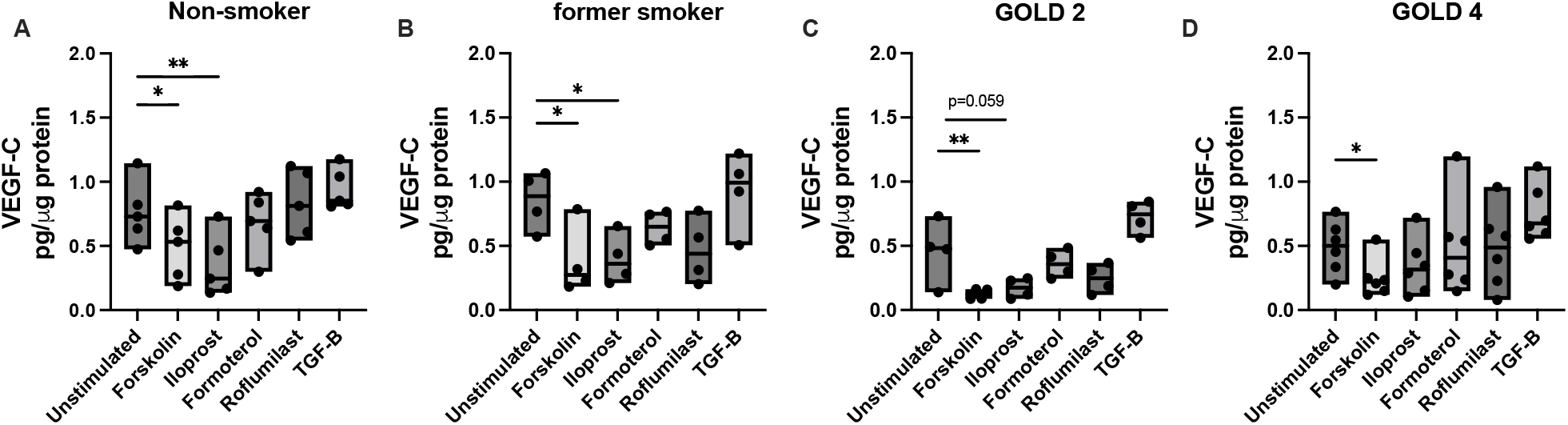
VEGF-C synthesis in lung fibroblasts. Effect of cAMP mediated compounds forskolin 1 μM, iloprost 1 μM, formoterol 1 μM and Roflumilast 1 μM, and transforming growth factor (TGF)-β1on VEGF-C release in primary distally derived fibroblasts from healthy lung donors (n=6, A), former smokers (n=4, B), patients with COPD GOLD 2 (n=4, C) and GOLD 4 (n=6, D). Data is presented as min max with individual values. Median values are represented as a line. ^*^P < 0.05; ^**^P < 0.01. Statistical analyses were performed with analysis of variance (ANOVA) on ranks followed by Dunn’s post hoc test to compare differences between and within lung fibroblasts form control subjects (non-smokers and ex-smokers) and COPD subjects (GOLD 2 and GOLD 4) after treatments compared to unstimulated fibroblasts.

There were no significant differences in VEGF-A at unstimulated conditions or after TGF-b stimuli between control subjects (n=10) and COPD patients (n=10). When subgrouped, ex-smokers (n=4) showed a significantly higher release of VEGF-A (p<0.05) compared to non-smokers (n=6), COPD GOLD 2 (n=4) or GOLD 4 (n=6). The release of VEGF-A was significantly increased after stimulation with TGF-b1 10 ng/mL in lung fibroblasts from both control subjects and patients with COPD (Fig 1.A-D). Treatment with different cAMP mediating compounds showed that iloprost and forskolin significantly increased the release of VEGF-A in fibroblasts from healthy lung donors (fig 2.A) and in COPD GOLD 4 (fig 2.D), whereas only iloprost significantly increased VEGF-A in COPD GOLD 2 (Fig 2.C) and roflumilast in GOLD 4 (p=0.045, Fig 2.D). No significant effects of the cAMP mediated drugs were observed in fibroblasts obtained from ex-smokers (Fig 2B).

**Figure 2.**
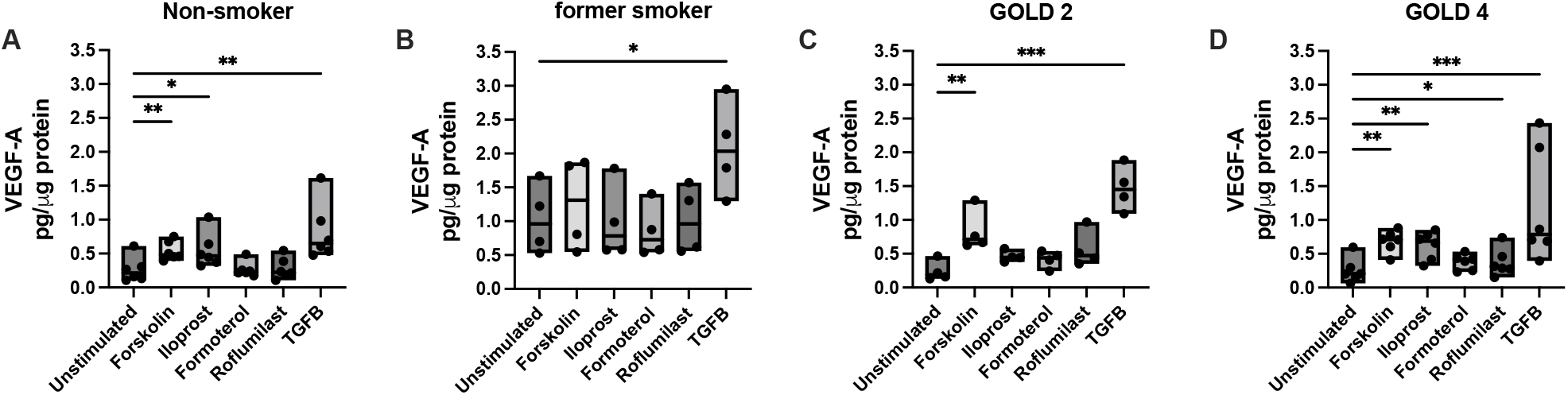
VEGF-A synthesis in lung fibroblasts. Effect of cAMP mediated compounds forskolin 1 μM, iloprost 1 μM, formoterol 1 μM and Roflumilast 1 μM, and transforming growth factor (TGF)-β 1on VEGF-A release in primary distally derived fibroblasts from healthy lung donors (n=6, A), former smokers (n=4, B), patients with COPD GOLD 2 (n=4, C) and GOLD 4 (n=6, D). Data is presented as min max with individual values. Median values are represented as a line. ^*^P < 0.05; ^**^P < 0.01; ^***^P < 0.001. Statistical analyses were performed with analysis of variance (ANOVA) on ranks followed by Dunn’s post hoc test to compare differences between and within lung fibroblasts form control subjects (non-smokers and ex-smokers) and COPD subjects (GOLD 2 and GOLD 4) after treatments compared to unstimulated fibroblasts.

**Figure 3.**
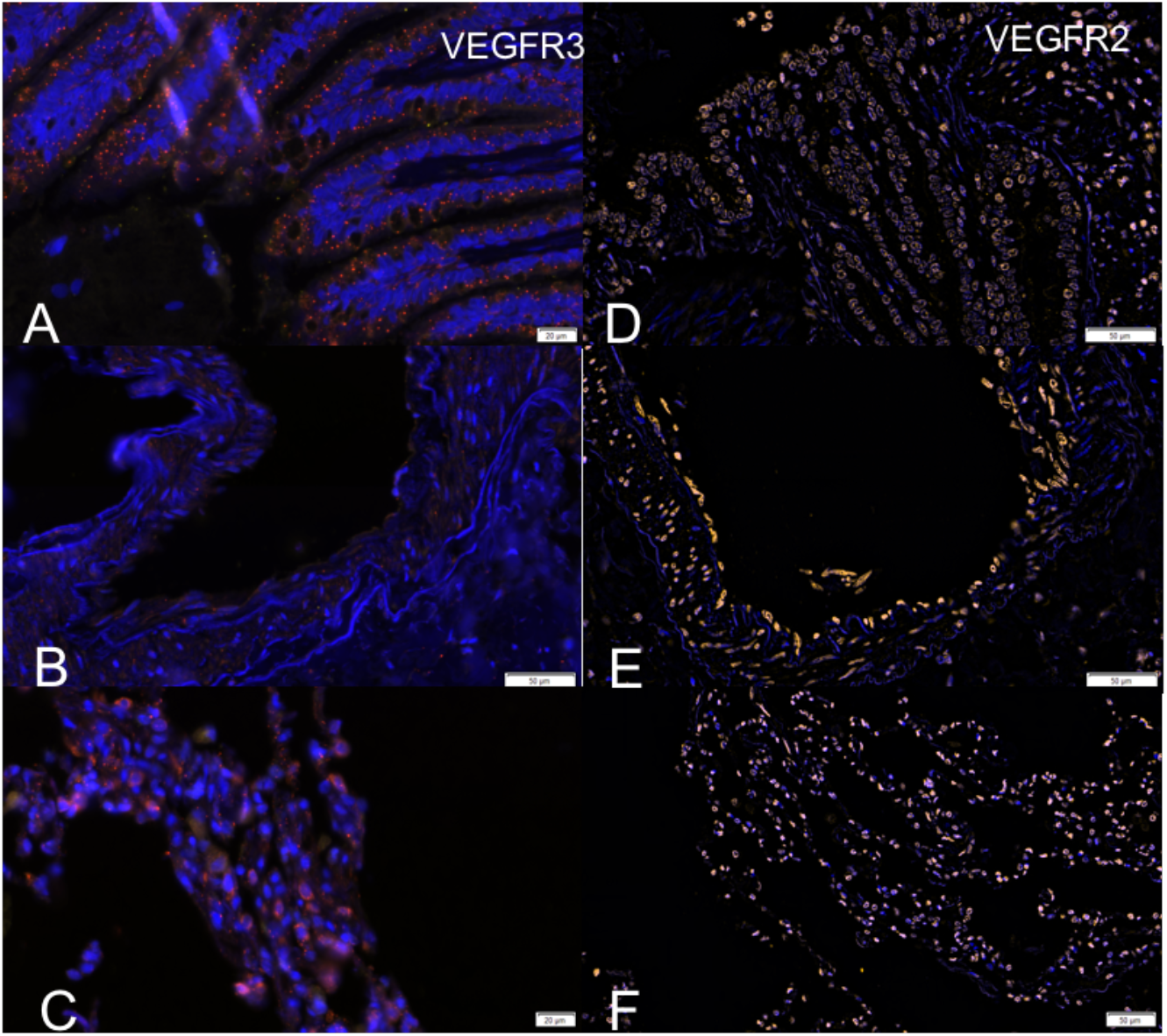
**A-F**: Immunofluorescence staining of different lung structures, the airway epithelium (A and D), endothelium (B and E) and parenchyma (C and F) in the distal lung from a healthy subject. A-C illustrates the expression of VEGFR3 in red, where a clear expression of VEGFR3 can be observed in the airway with a receptor-like morphology (A) and in the parenchyma with an expression both extra- and intracellularly (C) while almost no expression was observed in the endothelium (B). Co-staining with p4OH was observed but primarily located in the parenchyma. D-F illustrates the expression of VEGFR2 in yellow. VEGFR2 is less expressed in the bronchial epithelium (D) and more prominently expressed in the endothelium of vessels (E) and in the vasculature of the parenchyma (F) in the lung tissue.

### Expression of VEGFR2 and VEGFR3 in different compartments of the lung tissue

Immunofluorescence staining indicated that VEGFR3 was mainly expressed in bronchi and lung parenchyma (3A-C), whereas VEGFR2 was expressed mainly in capillaries in the parenchyma and in pulmonary vessels (3D-F) in lung tissue from a healthy individual.

Further staining of lung tissue from control subjects and patients with GOLD 2 and GOLD 4 confirmed expression of VEGFR3 in the bronchial epithelium. Immunofluorescence staining using the primary rabbit anti-human VEGFR3 antibody co-stained with the epithelial marker pan cytokeratin suggest that VEGFR3 expression increases progressively from healthy subjects (n=5) to COPD GOLD 2 (n=4) and GOLD 4 (n=5). Co-staining with the epithelial marker pan-cytokeratin confirmed that the staining of VEGFR3 was expressed on bronchial epithelial cells (fig 4).

**Figure 4.**
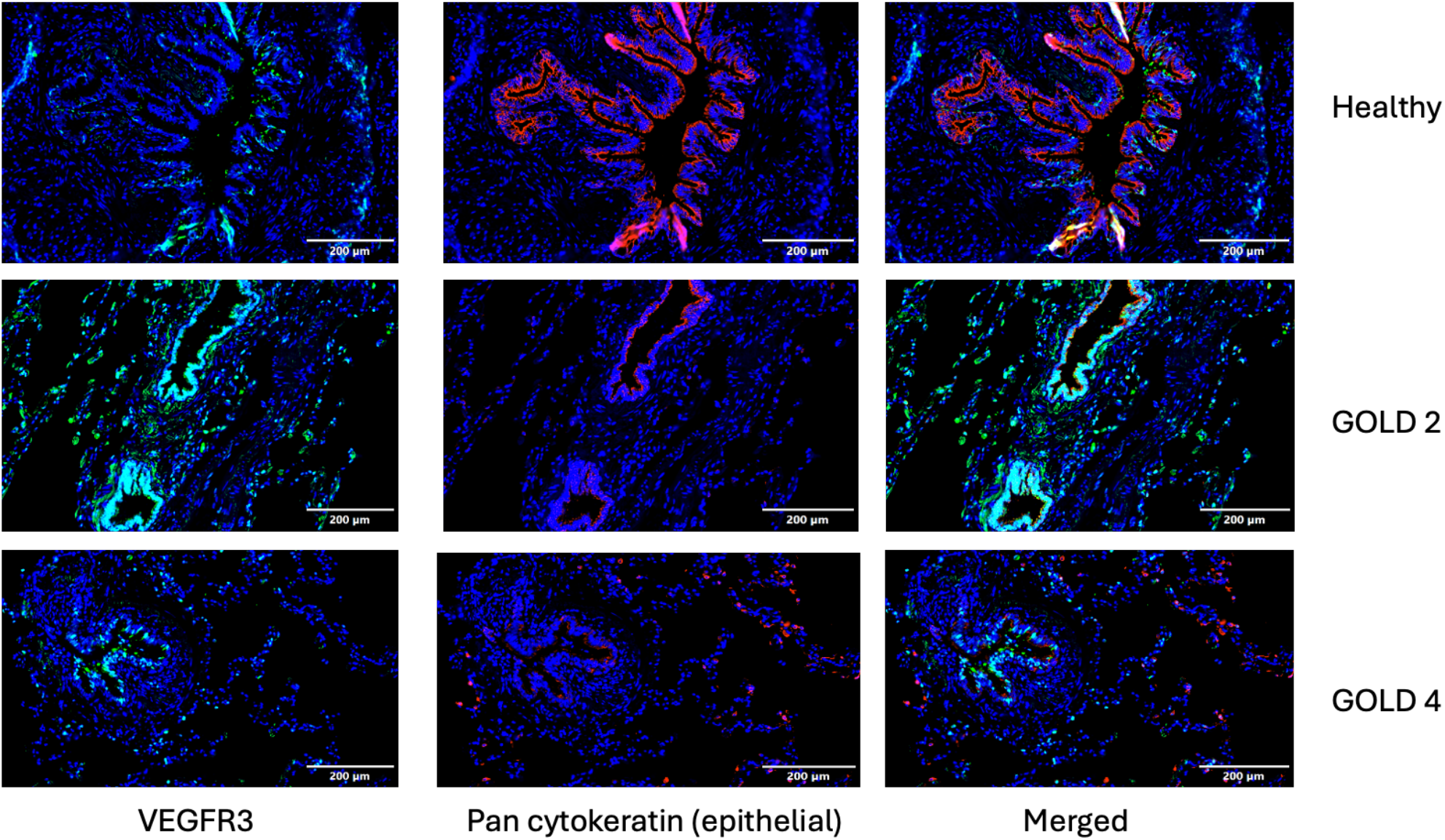
Immunofluorescence staining using the primary rabbit anti-human VEGFR3 antibody co-stained with the epithelial marker pan cytokeratin suggest that VEGFR3 expression increases from healthy to COPD GOLD 2) and COPD GOLD 4. Images of the airways from a healthy lung donor, a patient with COPD GOLD2 and a patient with COPD GOLD 4 are visualized.

Triple staining of lung tissue from a healthy lung donor (Fig 5A) and a patient with COPD GOLD 4 (Fig 5B) with vimentin and pan-cytokeratin indicated expression of VEGFR3 in the bronchial epithelium and on mesenchymal cells, as fibroblasts. Vimentin is also present in the endothelium layer but not in the bronchial epithelium, whereas pan-cytokeratin is co-expressed with VEGFR3 in the epithelium (Fig 5).

**Figure 5.**
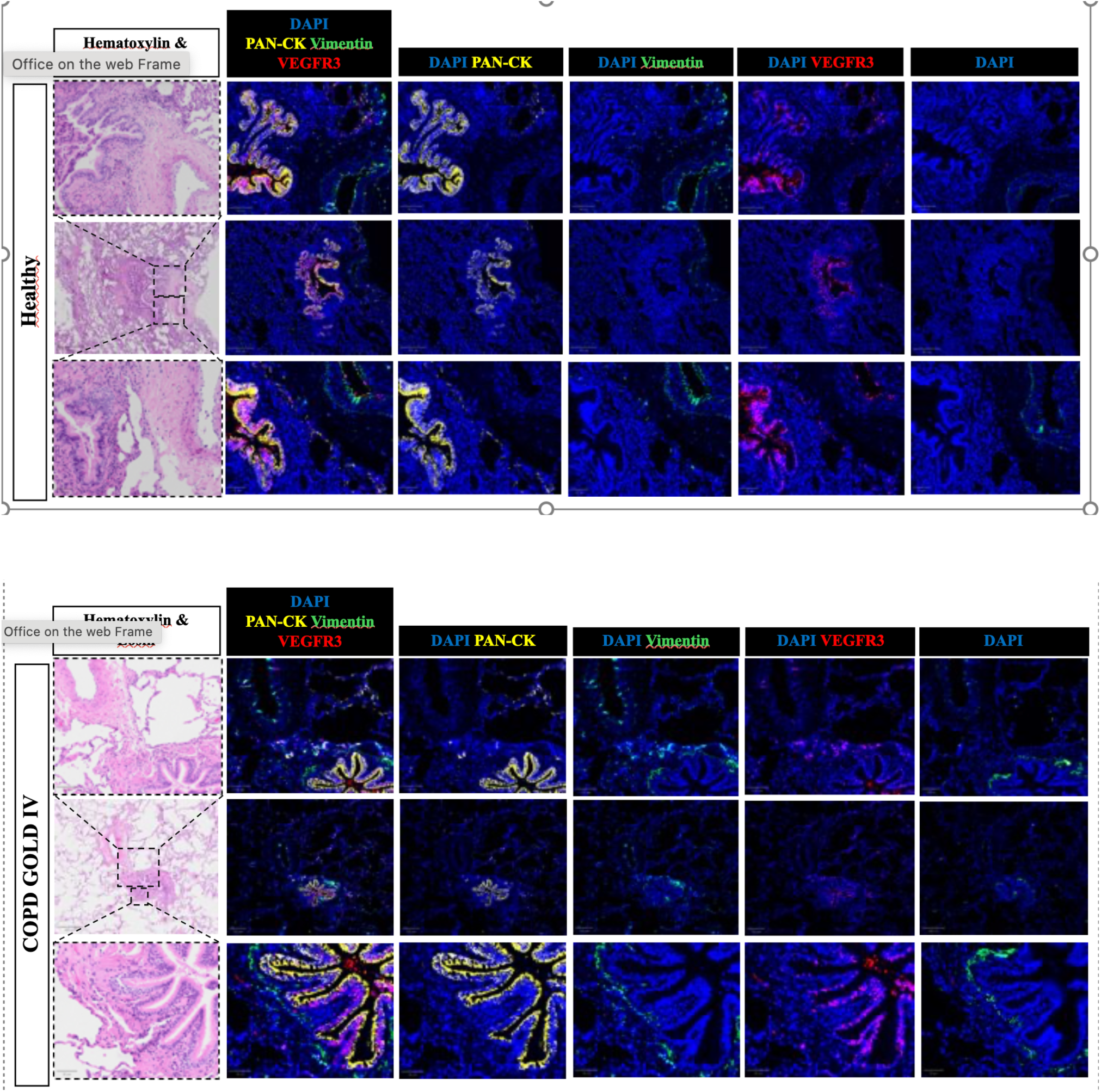
HE staining to visualize the lung tissue with vessel and bronchi. Immunofluorescence staining of lung tissue from a healthy control lung donor (top) and from a COPD GOLD 4 patient (bottom) showing staining of VEGFR3 (red) together with pan cytokeratin (yellow) and vimentin (green). VEGFR3 is present in the bronchi but not in the pulmonary vessel.

### Expression of VEGFR3 in lung fibroblasts and bronchial epithelial cells

To verify if the fibroblasts express VEGFR3 lysates from primary lung fibroblasts were analysed using western blot. Due to limited material three experiments with lysate from fibroblasts from healthy control subjects (non-smokers) and COPD GOLD 4 patients were evaluated for VEGFR expression and if the expression were altered after treatments with iloprost, formoterol or forskolin. The major precursor of VEGFR-3 matures to a 195 kDa fully glycosylated cell-surface receptor, which is then proteolytically cleaved into 125 kDa C-terminal fragment, and a shorter N-terminal fragment (20). Using a VEGFR-3 antibody that detects the bands at 175 kDa, 125 kDa, 80 kDa, we consistently observed across all blots a prominent ∼75 kDa protein band (Fig. 6A, 6C). Other faint bands were inconsistently observed and not reliably measurable. Truncated forms of VEGFR3 have been described in other cell types, suggesting that proteolytic processing may generate lower-molecular-weight fragments detectable by immunoblot (21). Fibroblasts displayed the ∼75 kDa band recognized by the anti-VEGFR3 antibody. Quantification of this band revealed no significant changes following treatment with either forskolin, iloprost or formoterol compared to untreated lung fibroblasts in either healthy control (A, C) or COPD fibroblasts (B, D).

**Figure 6.**
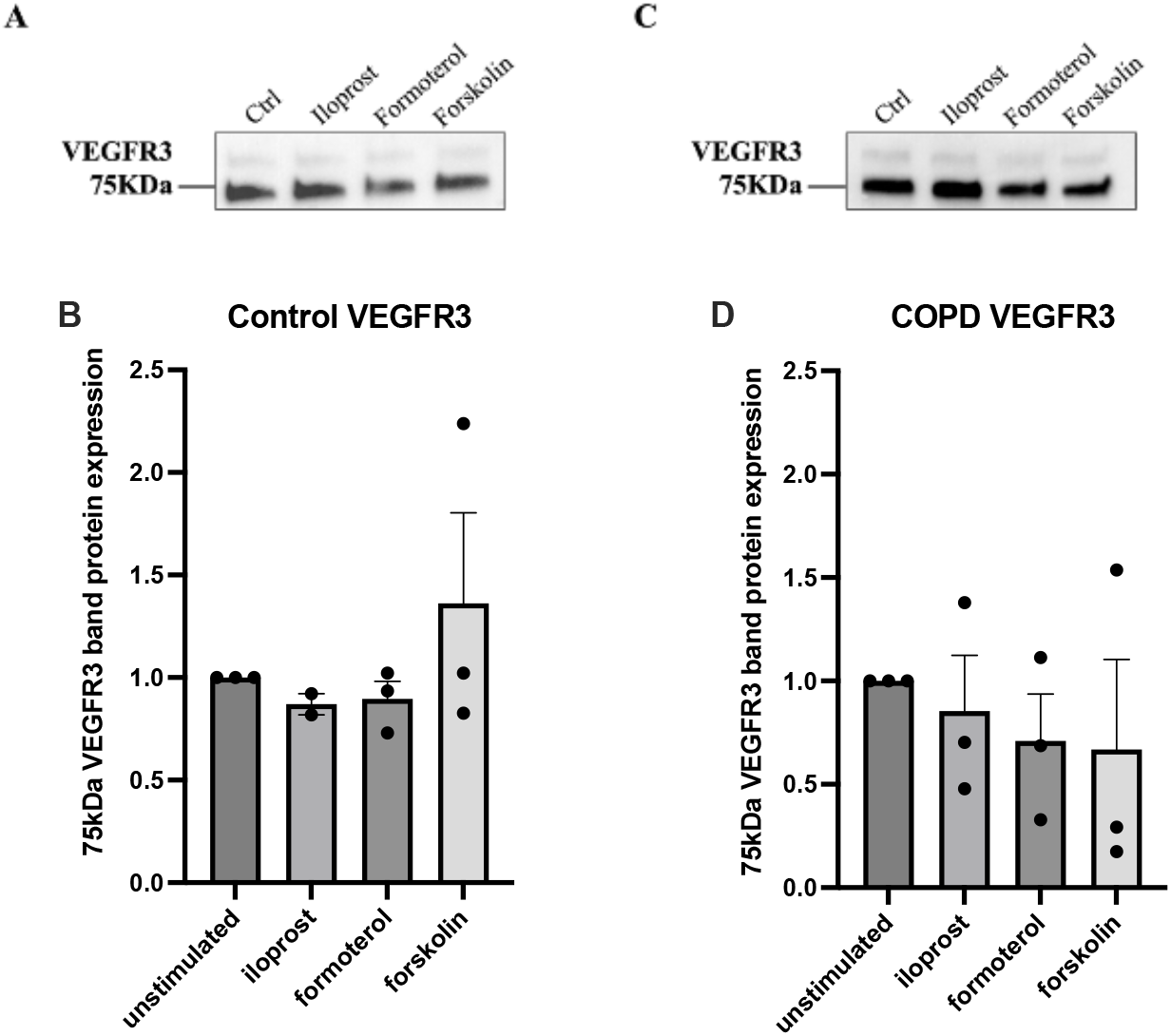
Immunoblot detection of VEGFR3 75 kDa fragment in lysates of fibroblast derived from healthy control subjects (A) and COPD GOLD 4 patients (C) treated for 24 h with iloprost, formoterol or forskolin. Bands are representative of one out of three separate experiments. Densitometric quantification of VEGFR3 expression in fibroblasts lysates from controls (B) and COPD patients (D). Data is shown as bars with individual values and SEM.

To further investigate the role of VEGF-C and the increased VEGFR3 expression in bronchi, studies with VEGF-C exposure were performed in bronchial epithelial cells using the cell line BEAS-2B and in human lung fibroblasts (HFL-1). Quantification of the 75 kDa protein band in lysate from BEAS-2B and HFL-1 cells, after stimulation with either 1 or 10 ng/mL of VEGF-C, that activates both to VEGFR2 and VEGFR3, or VEGF-C (Cys156Se), that selectively activates VEGFR3, or 10 ng/mL of TGF-β, did not show any statistically significant alterations compared to untreated control in expression of VEGFR3 (n=8) (Fig 7 A-D). There was however a trend towards increased expression of VEGFR3 (p=0.075) in HFL-1 after exposure to 1 ng/mL of VEGF-C (fig 7 B).

**Figure 7.**
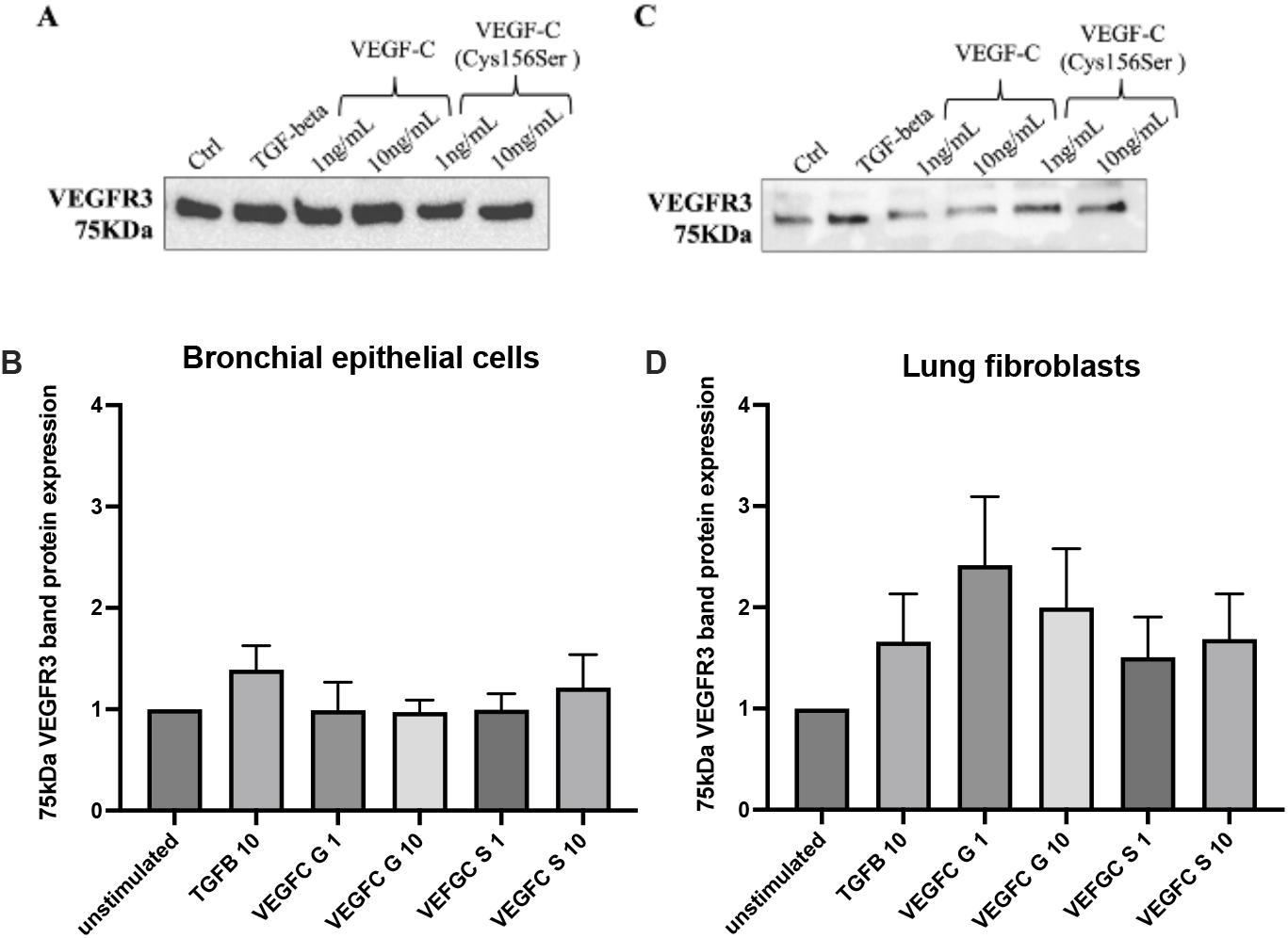
Immunoblot detection of VEGFR3 75 kDa fragment in lysates of bronchial epithelial cells BEAS2B (A, B) and human lung fibroblasts HFL-1 (C, D) treated for 24 h with 10 ng/mL of TGF-b1 and 1 ng/mL or 10 ng/mL of VEGF-C and VEGF-C (Cys156Ser). Bands are representative from one out of four separate experiments in BEAS-2B (A) and HFL-1 (C)). Densitometric quantification of VEGFR3 expression in lysates of BEAS-2B (B) and HFL-1 cells (D). Data were normalized against total protein amount. Data are expressed as mean ± SEM, n = 4 independent experiments with two biological replicates in each experiment. Statistical analysis was performed with One sample t test.

### Effect of VEGF-C stimulation on proliferative rate

We have previously shown that exposure to VEGF-A165 increased proliferative rate in HFL-1. The VEGF-C (Cys156Ser), selective ligand for VEGFR3, significantly reduced proliferative capacity after 48 hours of exposure both in HFL-1 and in BEAS-2B, whereas the unselective VEGF-C that acts on both VEGFR2 and VEGFR3 did not affect the proliferative rate compared to control (Fig 8.A and 8.B).

**Figure 8.**
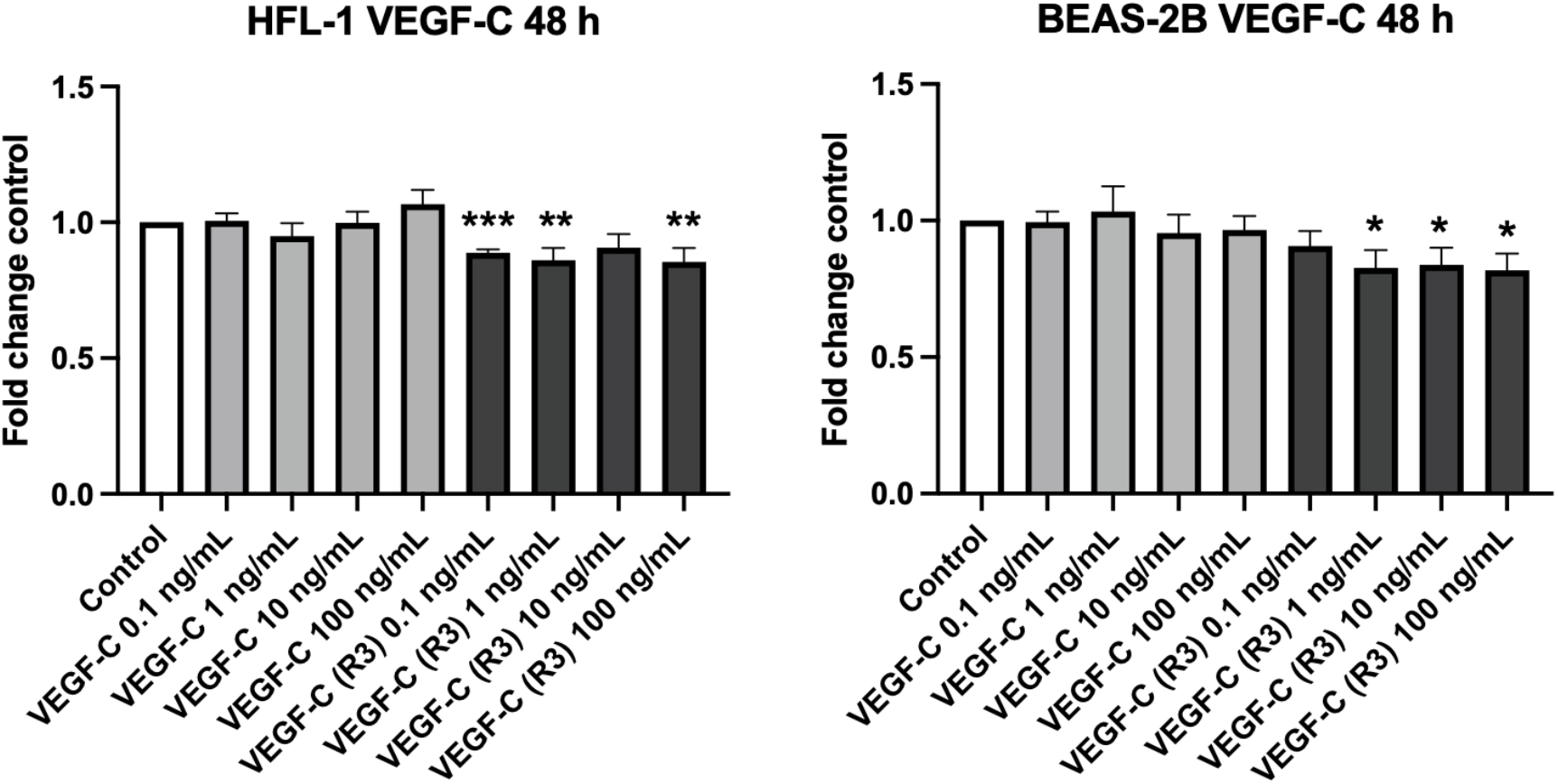
Proliferative capacity in human lung fibroblasts (HFL-1) (A) and bronchial epithelial cells (BEAS-2B) (B) treated with 1 ng/mL or 10 ng/mL of VEGF-C and VEGF-C (Cys156Ser) (VEGF-C (R3)). (n=4 independent experiments with 5 technical replicates in each. ^*^P < 0.05; ^**^P < 0.01; ^***^P < 0.001. Statistical analyses were performed with one-sample t-test compared to control.

## Discussion

Pulmonary vascular remodelling is commonly occurring in COPD. VEGF-A and its receptor VEGFR2 are known to be involved in angiogenesis and vascular remodelling in COPD (2, 3, 5), whereas VEGF-C and the receptor VEGFR3 are less investigated in COPD pathology. In this study we have focused on VEGF-C and its receptor VEGFR3 to further understand release and expression pattern in health and disease and how different cAMP generating therapies may affect the secretion of the VEGF isoforms VEGF-A and VEGF-C. The release of VEGF-A was significantly higher in lung fibroblasts obtained from former smokers compared to non-smokers and COPD patients, which has also been observed in a previous study on lung tissue from COPD patients, non-smoker and smoker subjects (2). In the present study, the release of VEGF-C was significantly lower in lung fibroblasts from COPD patients compared to controls. Profibrotic stimuli with TGF-β 10 ng/mL significantly increased the release of VEGF-A in both healthy and COPD-derived lung fibroblasts, whereas VEGF-C was not significantly altered after TGF-β stimuli, which is in line with our previous studies with primary human lung fibroblasts (13, 14). We have previously shown in lung fibroblasts that VEGF-A is induced by stimuli with either TGF-β or the prostacyclin analogue iloprost and that there were differences in the response to iloprost between fibroblasts from very severe COPD patients (GOLD 4) and control subjects (no GOLD 2 patients were included in the study) (14). Our obtained data showed that VEGF-A increased proliferation and migration rate and enhanced biglycan and perlecan synthesis in an autocrine fashion in human lung fibroblasts (HFL-1), which is of importance for tissue integrity in the lung and vascular remodelling processes (14). The cellular response to the vasodilator iloprost was however dysregulated in primary distally derived lung fibroblasts from patients with very severe COPD (GOLD stage 4) (15). Iloprost reduced collagen I synthesis but did not affect the collagen-associated proteoglycans decorin and and biglycan in fibroblasts from COPD patients, which could lead to improper collagen network fibrillogenesis and a more emphysematous lung structure in COPD patients (15). In this study, we further evaluated the effect of iloprost and other cAMP generating therapies; formoterol, roflumilast that are currently prescribed to COPD patients, and forskolin that directly activates adenylyl cyclase and cAMP signalling, on the effect of VEGF-A and VEGF-C synthesis. Our obtained data indicate that cAMP mediating compounds in different degrees alter VEGF-A and VEGF-C synthesis in lung fibroblasts obtained from patients with different disease progression and GOLD stages compared to control subjects, implicating a dysregulated signalling of VEGF-A and VEGF-C in COPD. Iloprost induced a significant increase in VEGF-A release in lung fibroblasts from non-smoking controls and COPD GOLD 4, as we previously have observed (14), whereas no significant effects on VEGF-A release were observed in fibroblasts from former smokers or COPD GOLD 2 from cAMP generating therapies. The adenylyl cyclase activator forskolin induced an increase of VEGF-A synthesis and a decrease in VEGF-C synthesis in both control and COPD fibroblasts. On the contrary, the release of VEGF-C was significantly altered after exposure to iloprost and forskolin in fibroblasts from former smokers, non-smokers and patients with GOLD 2, whereas only forskolin affected the synthesis of VEGF-C in COPD GOLD 4. This data implicates that there is an imbalance between VEGF-A and VEGF-C signalling in fibroblasts from ex-smokers and patients with COPD. VEGF-C is known to promote lymphangiogenesis via VEGFR3 but also angiogenesis via activation of VEGFR2 by forming a complex where VEGF-C and VEGF-A stimulate angiogenesis in a synergistic manner (22-25). Activation of VEGFR3 may further control VEGFR2 expression and VEGF signalling to prevent excessive vascular permeability (26, 27). Staining of VEGFR2 and VEGFR3 was performed in healthy lung donor tissue and in lung tissue from COPD GOLD 2 patients obtained after lung resection and after lung transplantation of very severe COPD patients. VEGFR2 was mainly expressed in the lung parenchyma and in pulmonary blood vessels, whereas VEGFR3 was expressed in the lung parenchyma and in the bronchial epithelium. It appeared as VEGFR3 was more expressed in the bronchi in COPD, but further studies are needed to confirm these findings. VEGFR3 is expressed in lymphatic endothelial cells but is also distributed in many different organs including the lung, except for the endothelium of pulmonary blood vessels (9, 28). In our previous study VEGF-A increased proliferative rate in HFL-1, whereas in the present study the selective VEGF-C ligand decreased the proliferative rate in both HFL-1 and BEAS-2B, highlighting the regulation of VEGF-A/C signalling and receptor activation (25).

In conclusion, our study highlights the regulatory signalling of VEGF-A and VEGF-C in lung fibroblasts during homeostasis and indicates that VEGF-A and VEGF-C release and response to cAMP generating drugs are dysfunctional in fibroblasts from ex-smokers and patients with COPD.

We have further confirmed that VEGFR3 is highly expressed in the bronchial epithelium in lung tissue from COPD patients but also some control subjects. Importantly, further studies are needed to understand the role of VEGF-C and VEGFR3 in ongoing remodelling processes and the role of vascular changes in disease progression with the goal to find relevant biomarkers and treatment strategies for patients with COPD.

## Acknowledgements

This study was funded by the Swedish Heart-Lung foundation [20230453, 20240890, 20240890], the Swedish Research Council [2020/01375], Lund University Medical Faculty, Region Skane (ALF) 2022 − 0130, Birgit and Sven Håkan Ohlsson Foundation, the Crafoord foundation and Alfred Österlund foundation.

